# Novel Mutations in NSP1 and PLPro of SARS-CoV-2 NIB-1 Genome Mount for Effective Therapeutics

**DOI:** 10.1101/2020.12.02.408229

**Authors:** Mohammad Uzzal Hossain, Arittra Bhattacharjee, Md. Tabassum Hossain Emon, Zeshan Mahmud Chowdhury, Md. Golam Mosaib, Md. Moniruzzaman, Md. Hadisur Rahman, Md. Nazrul Islam, Irfan Ahmed, Md. Ruhul Amin, Asif Rashed, Keshob Chandra Das, Chaman Ara Keya, Md. Salimullah

## Abstract

Severe Acute Respiratory Syndrome Coronavirus-2 (SARS-CoV-2), the etiologic agent of Coronavirus Disease-2019 (COVID-19), is rapidly accumulating new mutations. Analysis of these mutations is necessary for gaining knowledge regarding different aspects of therapeutic development. Recently, we have reported a Sanger method based genome sequence of a viral isolate named SARS-CoV-2 NIB-1, circulating in Bangladesh. The genome has four novel mutations in V121D, V843F, A889V and G1691C positions. V121D substitution has the potential to destabilize the Non-Structural Protein (NSP-1) which inactivates the type-1 Interferon-induced antiviral system hence this mutant could be the basis of attenuated vaccines against SARS-CoV-V843F, A889V and G1691C are all located in NSP3. G1691C can decrease the flexibility of the protein while V843F and A889V changed the binding pattern of SARS-CoV-2 Papain-Like protease (PLPro) inhibitor GRL0617. V843F PLPro showed reduced affinity for Interferon Stimulating Gene-15 (ISG-15) protein whereas V843F+A889V double mutants exhibited the same binding affinity as wild type PLPro. Here, V843F is a conserved position of PLPro that damaged the structure but A889V, a less conserved residue, most probably neutralized that damage. Mutants of NSP1 could provide attenuated vaccines against coronavirus. Also, these mutations of PLPro could be targeted to develop anti-SARS therapeutics.

## 1. Introduction

Coronavirus Disease-2019 (COVID-19) is now an irrepressible pandemic that continuously threatening humanity since its causative agent Severe Acute Respiratory Syndrome Coronavirus-2 (SARS-CoV-2) is rapidly changing and adapting in different environments [1]. At the beginning of the COVID-19 outbreak, this viral infection caused flu-like symptoms such as cough, fever, and trouble breathing [2]. However, this acellular pathogen is constantly evolving with novel mutations thus viral diarrhea, cardiovascular injury, strokes, psychosis, dementia and Kawasaki like syndrome are becoming more frequent in COVID-19 cases [3–5]. According to the World Health Organization (WHO), this constant evolution in different selection pressures are letting the virus to be airborne [6]. Proper analysis of these genomic mutations can help to decipher issues regarding viral transmission, epidemiology, pathogenicity and therapeutics.

Coronaviruses (CoVs) are enveloped viruses that belong to the *Coronaviridae* family and *Orthocoronavirinae* subfamily [7, 8]. SARS-CoV-1 and Middle East Respiratory Syndrome (MERS-CoV) belong to the *Betacoronavirus* genus of *Orthocoronavirinae* subfamily [7, 8]. These pathogens were the causative agents of SARS and MERS epidemics. SARS-CoV-2, a member of *Betacoronavirus,* shares nearly 82.30% sequence similarity with SARS-CoV-1 while showing strikingly less similarity with MERS (28%) [9].

SARS-CoV-2 has a 29.9 kb single-stranded (+) positive RNA genome with 11 genes. The virus also contains two Untranslated Regions (UTRs) in 5’ and 3’ terminal. Nearly 67% of the genome consist of Orf1ab polyprotein gene. The rest of the genome constructs structural proteins e.g., Surface (S), Envelope (E), Membrane (M), and Nucleocapsid (N) proteins [10]. According to the recent update of Global initiative on sharing all influenza data (GISAID) database, with particular prevalent mutations, the virus has made six distinct clades such as S, L, V, G, GR and GH [11]. Among them, GR is the largest and predominates in South America, Europe and Asia (gsaid.org). The members of this clade contain S-D614G and N-G204R mutations. Currently, GR is relatively less prevalent in North America. However, recently we have sequenced a GR type viral isolate SARS-CoV-2/human/BGD/NIB_01/2020 (SARS-CoV-2 NIB-1) (NCBI GenBank Accession: MT509958, GISAID Accession ID: EPI_ISL_447904) from a young female COVID-19 patient of Bangladesh [12]. According to our report, the viral isolate shares a common ancestor with three North American SARS-CoV-2 isolates that were collected from Washington, United States of America (USA). Additionally, our isolate showed longer branch length or more genomic mutations than its closely related members [12]. When we aligned the genome with the reference genome (Accession: NC_045512.2), we found 11 mutations of which 4 of them were new. Among them, three were found in Papain-Like Protease (PLPro) and C-terminal domain of Non-structural Protein (NSP)-3 and another one was located in NSP1 (Table: I).

Several studies have demonstrated that NSP1 and the PLPro of NSP3 from SARS-Cov-1 and 2 play a critical role to inhibit the type 1 interferon (IFN) response in the host cell [13]. Moreover, PLPro cleaves off Interferon Stimulating Gene-15 (ISG15) protein, therefore, the host cell cannot execute antiviral signals properly [14, 15].

In this study, we have analyzed the effect of these novel mutations on the respective protein structures. We have shown that some of these mutations have the potential to destabilize the structure of NSP1 and the C-terminal domain of NSP3. We also explored how the loss and gain of Valine in V843F and A889V mutations respectively affect the binding of protease inhibitor GRL0617 and ISG15 for their implication in new therapeutics development.

## 2. Methods

### 2.1. Genome retrieval and identification of the mutations

The genome of SARS-CoV-2 NIB-1 was collected from the National Center for Biotechnology Information (NCBI) (GenBank accession: MT509958.1). The genome was aligned with the reference genome by NCBI Nucleotide Basic Local Alignment Search Tools (BLASTN) to identify the mutations in the Untranslated Regions (UTRs) [16]. The Non-synonymous mutations were collected from GISAID CoVsurver (gisaid.org).

### 2.2. Effect of the mutations on the proteins

Mutations that affect the structural stability of the proteins can behave differently than the wild type one [17, 18]. To assess the effect of the novel mutations, MUpro, Protein Variation Effect Analyzer (PROVEAN) and HOPE were employed [19–21]. Afterward, PLPro was taken for further analysis due to the availability of structural and molecular mechanisms on this protein.

### 2.3. *In silico* mutagenesis, molecular modeling and refinement of PLPro

To observe the mutational effect of the protein at three dimensional (3D) level, the crystal structure of PLPro was collected from Research Collaboratory for Structural Bioinformatics (RCSB) Protein Data Bank (PDB) (www.rcbs.org) [22]. The chain A of PLPro (PDB ID: 6W9C) was preserved via BIOVIA Discovery Studio (https://www.3ds.com/). Heterogeneous atoms were also removed. Mutated protein sequences went under homology modeling by SWISS-MODEL using chain A of 6W9C as a template [23]. The generated structures were energetically minimized by 3D Refine and then again refined by Galaxy Refine [24, 25].

### 2.4. Structural assessment of mutant proteins

The structural quality of the mutant proteins was evaluated. To execute this step, RAMPAGE and SWISS-MODEL structure assessment were employed [23, 26].

### 2.5. Inhibitor binding analysis against the wild type and mutant PLPros

GRL0617 or 5-Amino-2-methyl-N-[(R)-1-(1-naphthyl) ethyl] benzamide (PubChem CID: 24941262) is a PLPro inhibitor with a relatively low level of cytotoxicity [27]. Hence, this naphthalene based small molecule has the potential to be a prospective antiviral drug. Here, we analyzed the interactions between GRL0617 and PLPro by AutoDock Vina using our previously reported methodology for exploring drug-receptor interactions [28, 29]. In short, the GRL0616 was taken as ligand and the wild type and mutant proteins were used as receptors. The active site of the palm domain in PLPro was enclosed with Grid Box [30]. (Parameters are given in Supplementary file:1). The molecular interactions between GRL0617-PLPro were visualized by UCSF Chimera [31], BIOVIA Discovery Studio and PyMOL Molecular Graphics System, Version 2.3.3 Schrödinger, LLC.

### 2.6. Analysis of ISG15-PLPro interactions

ISG15 C-terminal domain that interacts with PLPro was collected from PDB ID: 6XA9. As a ligand/ substrate, this protein was docked against the PLPro receptors by GalaxyTongDock_A [32]. Model 1 with the highest cluster size and docking score was further analyzed by PRODIGY (PROtein binDIng enerGY prediction) to determine the binding affinity of the protein complexes at 25° C temperature [33].

## 3. Results

The overall scheme of the present study has been described in **Figure 1**

**Figure 1:**
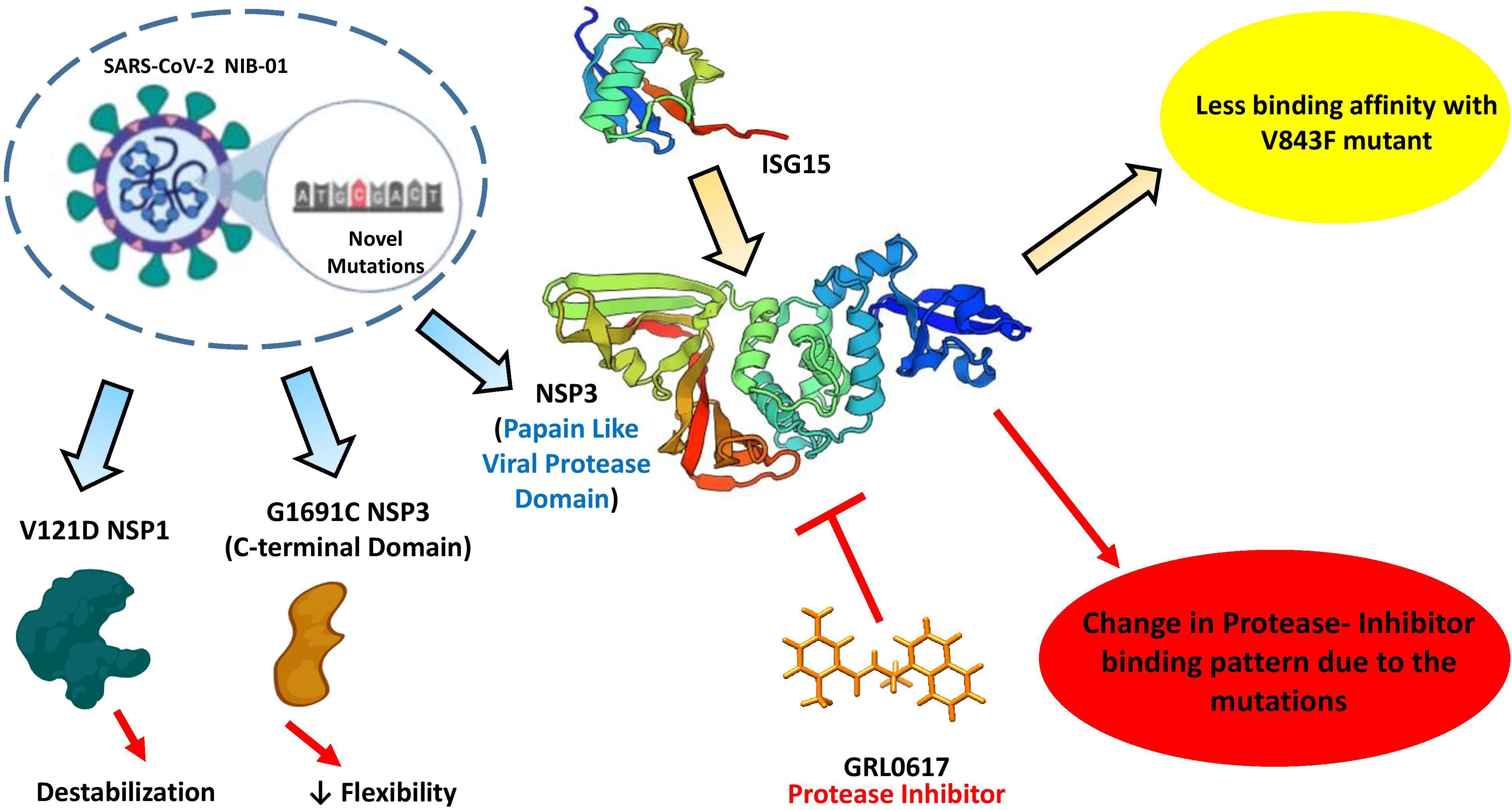
Graphical representation of the whole work. Some of the parts of this figure were generated with the help of BioRender (https://biorender.com/).

### 3.1. Transversion mutations are predominant in SARS-CoV-2 NIB-1

SARS-CoV-2 NIB-1 is a member of the GR clade (Table 1). The genome contains 11 mutations in total. Among them, 6 were transversion mutations and 5 were transition mutations. According to GISAID, 7 of these mutations were reported previously and 4 of them were first identified in SARS-CoV-2 NIB-1 isolate. In these 7 mutations, 4 were transition mutations; Three Purine

**Table 1:**
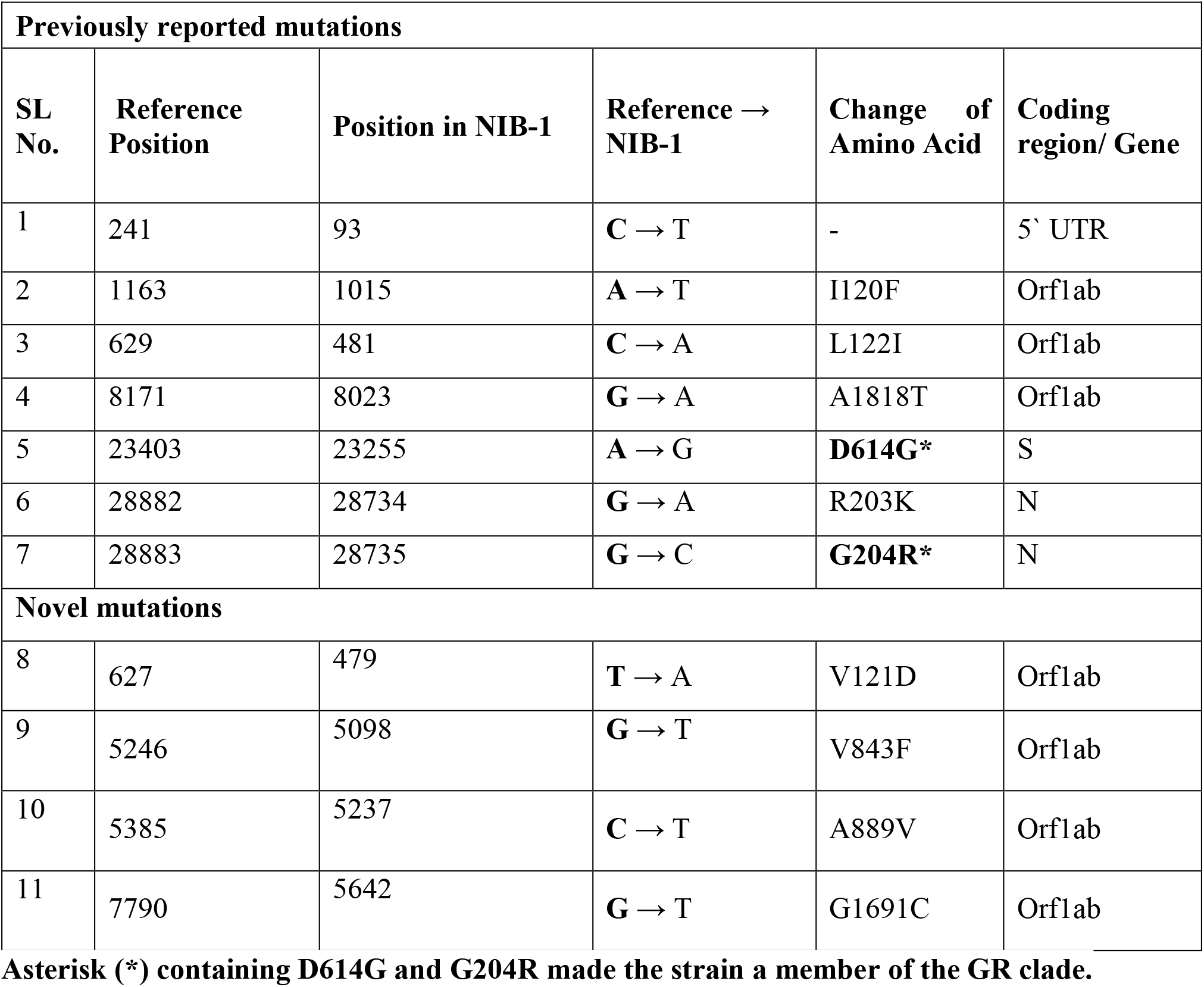
Mutations in SARS-CoV-2 NIB-1.

→Purine transitions (Table 1; Serial # 4-6) and one Pyrimidine →Pyrimidine transition (Table 1; Serial # 1). The other three mutations were transversion mutations: Two Purine → Pyrimidine transversions (Table 1; Serial # 2, 3) and one Pyrimidine →Purine transversion (Table 1; Serial # 7). The 4 novel mutations were located in the Orf1b gene. Among them, only A889V substitution occurred due to C → T (Pyrimidine → Pyrimidine) transition. The other 3 mutations were transversion substitutions; Two Purine → Pyrimidine transversions (Table 1; Serial # 9, 11), one Pyrimidine →Purine transversion.

### 3.2. Novel mutations have the potential to alter the functions and structures of protein

According to the MUPro, V121D, V843F and G1691C destabilize the proteins (Table 2). Moreover, PROVEAN determined that the V121D and G1691C substitutions are deleterious for its biological functions. HOPE demonstrated that most of the mutant amino acids are located in a conserved region and they are bigger than the wild type residues. They have the potential to abolish the protein functions. In contrast, only A889V mutant PLPro can increase protein stability (Table 2).

**Table 2:** Effect of the novel mutations in protein stability, function and structure.

| **SL No.** | **Change of Amino Acid** | **Protein Stability (MUPro)** | **Protein Function (PROVEAN; cutoff= -2.5)** | **Protein Structural Properties (HOPE)** |
| --- | --- | --- | --- | --- |
| 1 | V121D | DECREASE | Deleterious | **Change of Amino Acid :**<br>NEUTRAL→ NEGATIVE<br>**Change in Structural Stability:** Slight destabilization of PLPro<br>**Level of Conservation:**<br>Highly Conserved |
| 2 | V843F | DECREASE | Neutral | **Change of Amino Acid Size:**<br>Mutant residue size is bigger<br>**Change in structural stability:**<br>Disturbing the domain and might abolish its function<br>**Level of Conservation:**<br>Highly conserved |
| 3 | A889V | INCREASE | Neutral | **Change of Amino Acid Size:** |
|  |  |  |  | Mutant residue size is bigger.<br>**Change in Structural Stability:**<br>Disturbing the domain and might abolish its function<br>**Level of Conservation:**<br>Located near a highly conserved position, this mutation might occur without damaging the protein. |
| 4 | G1691C | DECREASE | Deleterious | **Change of Amino Acid Size:**<br>Mutant residue size is bigger and more hydrophobic.<br>**Change in Structural Stability:**<br>The wild-type residue is a glycine that might be necessary to give the protein essential flexibility. Mutation of this glycine might abolish the function of the protein.<br>**Level of Conservation:** |
|  |  |  |  | The mutant residue is located near a highly conserved position. The mutant and wild-type residue are not very similar. Based on this conservation information this mutation is probably damaging to the protein. |

### 3.3. Interaction patterns of GRL0617-PLPro altered in mutant proteases

The interactions between the inhibitor GRL0617 and PLPro were identified via molecular docking simulation. Before molecular docking, the quality of the receptors was evaluated by analyzing Ramachandran plots and MolProbity scores. Ramachandran plots showed that the mutant PLPros have 98.1-98.4% amino acids in favored regions (~98% expected). The MolProbity score of the wild type crystal structure was 2.55 whereas the mutants have MolProbity scores between 1.16-1.32 (Supplementary file 3). Therefore, after the homology modeling, the quality of the simulated structures were satisfactory for further analysis [26, 34].

V843F PLPro (−6.7 kcal/mol) and A889V PLPro (−6.9 kcal/mol) showed more binding affinity scores than the wild type one (−6.6 kcal/mol) (Table 3). However, V843F+ A889V PLPro showed slightly less affinity (−6.2 kcal/mol).

**Table 3:** Interaction between wild type and mutant SARS-CoV-2 Papain-Like viral protease with GRL0617 and ISG15.

| SARS-CoV-2 Papain-Like viral protease and GRL0617 interaction patterns |  |  |  |  |
| --- | --- | --- | --- | --- |
| Receptor | Binding affinity score (kcal/mol) | Participating amino acids in interaction |  |  |
| Wild type | -6.6 | ASP164, ARG166, GLU167, *SER170*, PRO248, TYR264 |  |  |
| V843F | -6.7 | ASP76, ARG82,*CYS155*,TYR171 |  |  |
| A889V | -6.9 | ASP76, ARG82, *CYS155*, TYR171 |  |  |
| V843F+<br>A889V | -6.2 | ASP76, ARG82, *THR75*, PRO77 |  |  |
| SARS-CoV-2 Papain-Like viral protease and ISG15 interaction patterns |  |  |  |  |
| Receptor | TongDock_A score | Cluster size | Binding affinity score (kcal/mol) | Dissociation constant (K<sub>d</sub>) (M) |
| Wild type | 915.278 | 41 | -15.4 | 4.8E-12 |
| V843F | 720.354 | 23 | -11.1 | 7.2E-09 |
| A889V | 1019.921 | 31 | -15.4 | 4.7E-12 |
| V843F+<br>A889V | 968.296 | 28 | -15.5 | 4.2E-12 |

Although the binding affinity did not change significantly due to the mutations, the inhibitor binding site changed and hence the interacting amino acids (Table 3). The wild type receptor interacted with ASP164, ARG166, GLU167, SER170, PRO248, TYR264 whereas ASP76, ARG82 were common interacting residues for mutant type receptors. Moreover, residues that participated in hydrogen bonding in Wild type single mutants and double mutants were not the same. V843F and A889V single mutants have the same residue CYS155 which interacted with the inhibitor through Hydrogen bonding.

### 3.4. V843F mutation significantly reduced the affinity of PLPro toward ISG15

SARS-CoV-2 PLPros cleaves off ISG15 that is essential for type 1 IFN mediated antiviral mechanisms [14]. The wild type, A889V and V843F+ A889V mutants showed nearly the same binding affinity (−15.4 to −15.5 kcal/mol) toward the C terminal domain of ISG15. However, V843F mutant exhibited a significantly lower docking score and binding affinity (−11.1 kcal/mol) with higher dissociation constant (Table 3).

## 4. Discussions

SARS-CoV-2 is a zoonotic pathogen that initiated the current global pandemic COVID-19 [2]. Since SARS-CoV-2 is a novel coronavirus and mostly resemble with SARS-CoV-1 (80% sequence similarity), we therefore focused on SARS-CoV-1 related reports to understand the results of the mutations [35]. However, we also explored other well studied *betacoronavirus* members such as Middle East Respiratory Syndrome (MERS-CoV), Human Coronavirus OC43, Mouse Hepatitis Coronavirus (MHV) to comprehend different genomic and proteomic functions of SARS-CoV-2.

Since the report of reference genome SARS-CoV-2 from Wuhan in December 2019, the deadly pathogen has spread throughout the world and produced numerous genetic variants with six predominant clades (www.gisaid.org). The GISAID CoVsurver analysis revealed that our previously reported isolate ‘SARS-CoV-2 NIB-1’ is a member of GR clade which has been mutated 10 times in the coding regions [12]. The NCBI BLASTN unveiled another mutation in 5’ UTR region. Interestingly, among these mutations, 54% occurred due to transversion mutations (Table 1) although species are usually biased to transition mutations [36]. Transversion mutations are detrimental for RNA viruses e.g. Influenza A and Human Immunodeficiency Viruses (HIV) since it can radically change the amino acids [37].

The novel mutations V121D, V843F and G1691C occurred in SARS-CoV-2 NIB-1 isolate due to transversion substitutions have the potential to destabilize or alter the protein structure and function (Table 2). Especially, V121D, V843F position are highly conserved in the protein and G1691C might reduce the essential flexibility of NSP3 (Table 2). Amino acid substitution in V843F and G1691C took place due to G → T transversion that was possibly introduced by Oxo-guanine generated from reactive oxygen species (ROS) [37, 38]. Conversely, C → T transition, the most frequent transition of SARS-CoV-2, increased the structural stability of PLPro (Table 2) [38]. Besides, the C **→** T transition in the 5’ UTR region might interfere with the function of N and NSP1 protein [39].

SARS-CoV-2 NIB-1 has V121D and L122I amino acid substitutions in the NSP1. SARS-CoV-2 NSP1 has 84.4% similarity with SARS-CoV-1 NSP1 [40]. NSP1 is a potent virulence factor of SARS-CoV-1that reduces the host mRNA expression by binding with the host 40S ribosome to inhibit the translation and specifically accelerate host mRNA degradation keeping the viral mRNA intact [40, 41]. NSP1 also disrupts the activation of IFN-dependent antiviral signaling pathways and represses the expression of the innate immune response genes such as type I IFN, ISG56 and ISG15 [13, 42]. That IFN and ISG15 gene-mediated antiviral pathways are crucial in host defense mechanisms against SARS-CoV-1 & 2 were shown by the suppression of their multiplication through *in vitro* type-1 IFN treatment [43, 44]. An attenuated mutant of NSP1 in SARS-CoV-1 can replicate as efficiently as wild-type strains with an intact IFN response [45]. Moreover, mutant NSP1 of MHV exhibited an adequate amount of cytotoxic T cells that protected the mice against further viral infections [46]. Hence, there is a higher possibility that a suitable NSP1 mutant of SARS-CoV-2 would work as an attenuated vaccine for COVID-19. Through computational analysis, we have observed that V121D and L122I mutant might destabilize the structure of NSP1 (Table 2 & Supplementary: 2) yet with these two mutations, the virus caused mild fever, cough and throat congestion in a young female patient [12]. Therefore, V121D and L122I mutations along with 93^th^ C **→** T in 5’ UTR did not change the pathogenicity of SARS-CoV-2 significantly.

However, keeping in mind that the quick recovery of the patient within 10 days of the onset of symptoms without any notable care, it might be speculated that NSP1 with these mutations could be a potent candidate for the development of an attenuated vaccine against SARS-CoV-2.

Main protease and papain-like protease (PLPro) are two essential viral enzymes that are vital for polypeptide processing during viral maturation [40]. Therefore, inhibitors of these proteins could be a prospective antiviral drug [47]. SARS-CoV-1 PLPro has been targeted to develop inhibitors for decades [30]. SARS-CoV-1 & 2 PLPros have 4 domains (Figure 3) [27]. The active site of this enzyme resides in the thumb domain. PLPro interacts with the N and C terminal domain of ISG15 through Ubiquitin binding subsites 2 and 1 respectively [30]. This interaction finally cleaves off ISG15. Hence, type 1 IFN induced ISG15 antiviral response cannot function properly in host cells [14]. A recent study also demonstrated that when SARS-CoV-1 & 2 infected Vero and Calu3 2B4 cell lines are treated with type 1 INF, the SARS-CoV-2 showed more susceptibility to the treatment [43, 44]. Most possibly the IFN treatment enhanced the synthesis of ISG15 that ceased the viral replications in the cell lines. PLPro mediated pathogenesis can be halted with naphthalene based protease inhibitor GRL0617 since this ligand can make non-covalent bonds with SARS-CoV-2 PLPro [14, 27]. Here, V843F and A889V mutations were located in the PLPro domain of NSP3 protein. V843F, A889V single mutants and V843F+A889V double mutants changed the binding site of GRL0617. The inhibitor binds in the thumb domain of the wild type receptor whereas the mutant receptors interacted through the Ubiquitin-Like Domain (Figure 2). Whether these alterations in binding patterns reduce the efficacy of the ligand or not is yet to be explored.

**Figure 2:**
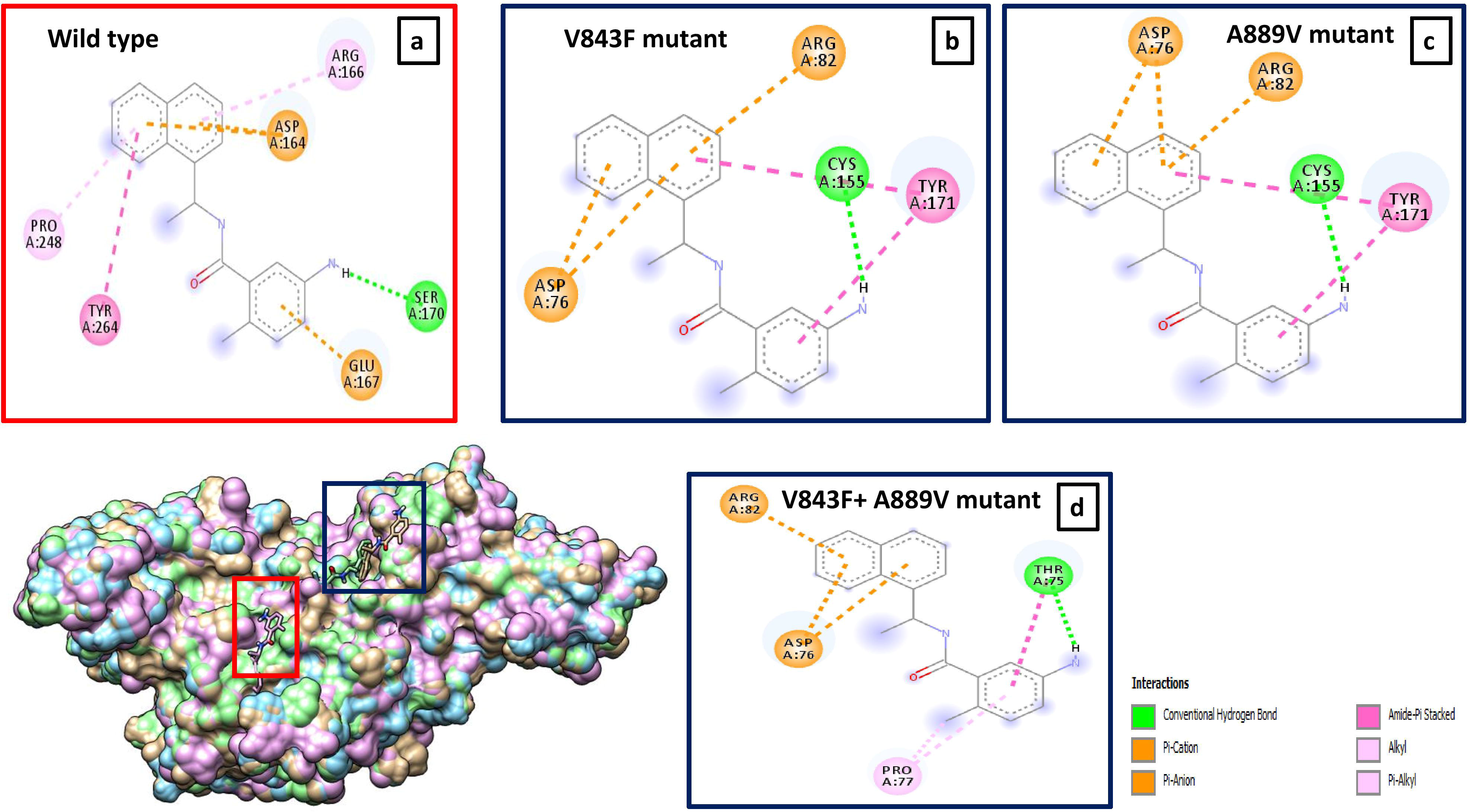
Interactions between GRL0617 inhibitor and superimposed SARS-CoV-2 wild type and mutant Papain-Like viral proteases (**a-d**). Interacting amino acid residues in wild type and PLPro mutant SARS-CoV-2. Inhibitor binding into the wild type and mutant protein. The red and blue boxes indicate the location and the interactions between ligand-receptors in wild and in mutant proteases respectively.

**Figure 3:**
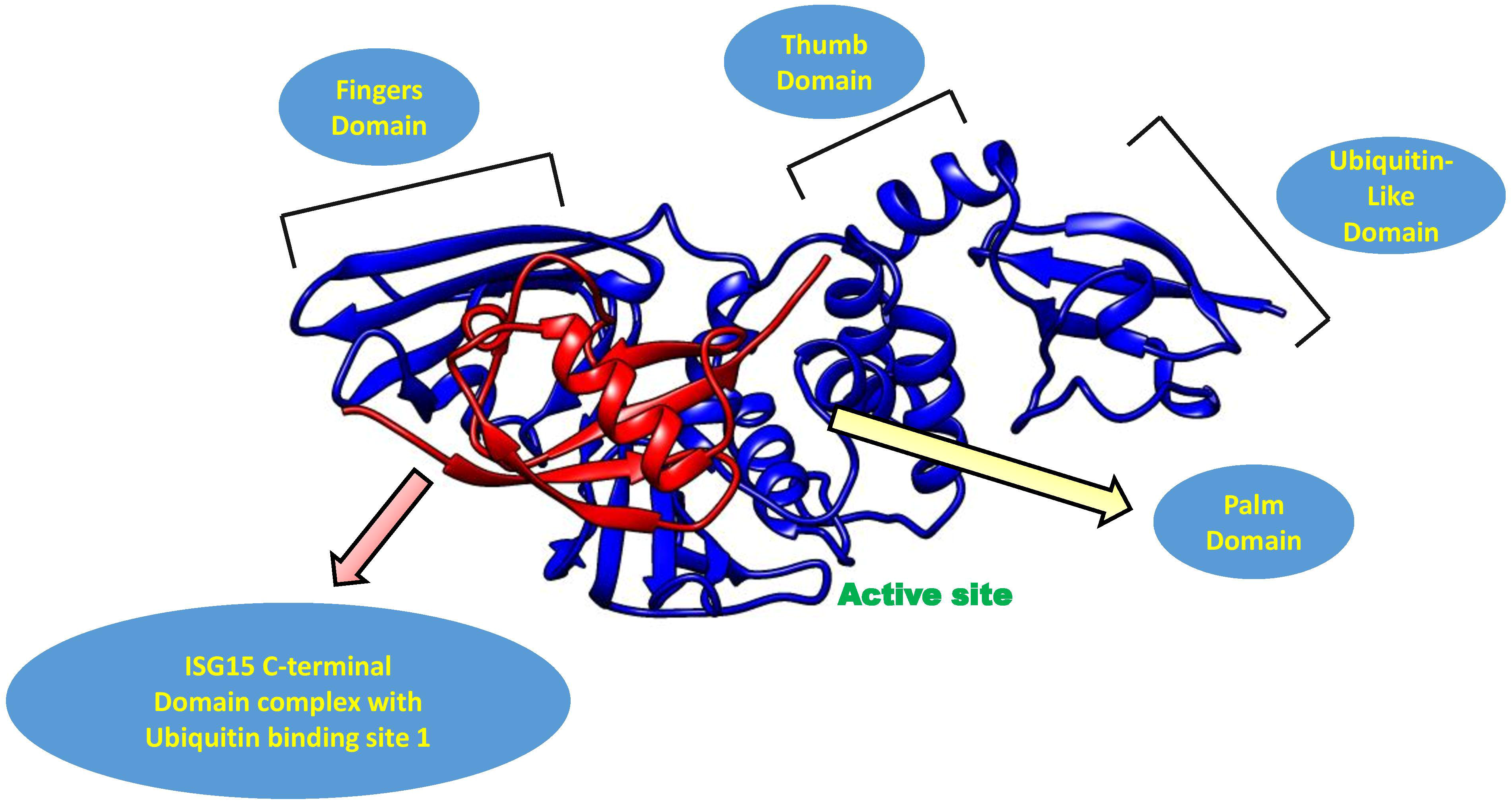
Interactions between the C-terminal domain of Interferon-Stimulated Gene 15 (ISG15) and SARS-CoV-2 wild type Papain-Like viral proteases. The red chain is the C-terminal domain of ISG 15. The blue chain represents the protease.

The mutant PLPros were also docked against the C-terminal domain of ISG-15. To execute this step, the ISG15 portion was separated from the PLPro-ISG15 C-terminal domain complex (PDB ID: 6XA9). Only the V843F single mutant showed reduced affinity toward the Ubiquitin binding subsite 1. Mutant A889V, V843F+A889V and Wild type PLPro did not show any difference when docked against ISG15. Therefore, V843F is a detrimental and destabilizing transversion mutation that might reduce the degradation of ISG15. It seems like A889V substitution protected the protease from the damage of V843F. Since A889V occurred due to the most common C → T transition, plausibly this mutation appeared first in the protein and then V843F took place due to ROS. Yet, further studies are needed to explore how this “yin and yang” incident affects the viral physiology. We hope our findings will give better insights during the development of attenuated vaccine and antiviral drugs.

## 5. Conclusions

This study focused on some novel mutations in the NSP1 and NSP3 proteins of SARS-CoV-2. NSP1 is a potent virulence factor of *Betacoronavirus* and hence a potential target for an attenuated vaccine. PLPro of SARS-like coronavirus has appeared as suitable targets for the development of anti-SARS therapeutics. In-depth, mutational studies on these proteins gave us better insights into the SARS-CoV-2 viral structure and pathogenicity for their fittingness as effective therapeutics candidates. In the future, wet lab research on the mutants will accelerate research and developments regarding attenuated vaccines and antiviral drugs.

## 6. Acknowledgments

The authors are grateful to the Ministry of Science and Technology, Bangladesh for the extensive support during this work.

